# Tree species and genetic diversity increase productivity via functional diversity and trophic feedbacks

**DOI:** 10.1101/2022.03.25.485785

**Authors:** Ting Tang, Naili Zhang, Franca J. Bongers, Michael Staab, Andreas Schuldt, Felix Fornoff, Hong Lin, Jeannine Cavender-Bares, Andrew Hipp, Shan Li, Yu Liang, Baocai Han, Alexandra-Maria Klein, Helge Bruelheide, Walter Durka, Bernhard Schmid, Keping Ma, Xiaojuan Liu

**Affiliations:** State Key Laboratory of Vegetation and Environmental Change, Institute of Botany, Chinese Academy of Sciences, 100093 Beijing, China; College of life sciences, University of Chinese Academy of Sciences, 100049 Beijing, China; College of Forestry, Beijing Forestry University, 100083 Beijing, China; Ecological Networks, Technical University Darmstadt, 64287 Darmstadt, Germany; Forest Nature Conservation, Georg-August-University Göttingen, 37077 Göttingen, Germany; Nature Conservation and Landscape Ecology, University of Freiburg, 79106 Freiburg, Germany; Institute of Applied Ecology, School of Food Science, Nanjing Xiaozhuang University, 211171 Nanjing, China; Department of Ecology, Evolution, and Behavior, University of Minnesota, 55108 St. Paul, USA; The Morton Arboretum, Lisle, 60532 IL, USA; State Key Laboratory of Systematic and Evolutionary Botany, Institute of Botany, Chinese Academy of Sciences, 100093 Beijing, China; Chair of Nature Conservation and Landscape Ecology, Faculty of Environment and Natural Resources, University of Freiburg, 79106 Freiburg, Germany; Institute of Biology/Geobotany and Botanical Garden, Martin Luther University Halle- Wittenberg, 06108 Halle, Germany; German Centre for Integrative Biodiversity Research (iDiv) Halle-Jena-Leipzig, 04103 Leipzig, Germany; Department of Community Ecology, Helmholtz Centre for Environmental Research–UFZ, 06120 Halle (Saale), Germany; Department of Geography, University of Zurich, CH-8057 Zurich, Switzerland

## Abstract

Addressing global biodiversity loss requires an expanded focus on multiple dimensions of biodiversity. While most studies have focused on the consequences of plant interspecific diversity, our mechanistic understanding of how the diversity within a given plant species (genetic diversity) affects plant productivity remains limited. Here, we use a tree species × genetic diversity experiment to disentangle the effects of species diversity and genetic diversity, and how they are related to tree functional diversity and trophic feedbacks. Tree species as well as genetic diversity increased tree productivity via increased tree functional diversity, reduced soil fungal diversity and marginally reduced herbivory. The effect of tree genetic diversity on productivity was partly different between tree species monocultures and mixtures: the functional diversity effect resulting from tree genetic diversity was only found in tree species monocultures, but the trophic feedbacks via herbivory were similar in species monocultures and mixtures. Given the complexity of interactions between tree species and genetic diversity, tree functional diversity and trophic feedbacks on productivity, we suggest that both tree species and genetic diversity should be considered in reforestation.

## Introduction

Biodiversity is essential for maintaining ecosystem functioning and nature’s contributions to people (Cardinale et al., 2012; Diaz et al., 2019). Ongoing biodiversity loss has received widespread concern from the international community (Ceballos et al., 2015). Expanding our research focus to multiple dimensions of biodiversity helps us to better predict the consequences of biodiversity loss and prioritize the different dimensions of biodiversity in conservation efforts (Cardinale et al., 2012). Whereas many studies related to biodiversity–ecosystem functioning (BEF) have focused on how interspecific diversity (i.e. the number of species) affects key ecosystem functions such as plant productivity (Hector et al., 1999; Huang et al., 2018; Tilman et al., 2001), relatively few have addressed effects of intraspecific diversity (such as genetic variation within a species). Furthermore, the effects of intraspecific diversity show an inconsistent picture: genetic diversity has promoted plant community productivity in herbaceous plant communities (Crutsinger et al., 2006; Kotowska et al., 2010) but not in forests (Bongers et al., 2020; Fischer et al., 2017). To get a better understanding of how genetic diversity influences plant productivity in forests and thereby could help guiding reforestation priorities we need to disentangle the underlying mechanisms.

Functional trait diversity, in short functional diversity, is expected to promote community productivity because different species or genotypes with diverse traits may use resources in complementary ways and then enhance the total utilization of resources in the whole community (Díaz & Cabido, 2001) (Fig. 1a). Thus, functional diversity, which has been primarily identified as the variation of plant functional traits among species, is often used to explain how plant species diversity impacts plant productivity (Cadotte et al., 2011; Díaz et al., 2007; Hillebrand & Matthiessen, 2009). Although genetic diversity within a species has been shown to explain substantial trait variation (Bongers et al., 2020), and intraspecific trait variation may have strong effects on plant productivity (Des Roches et al., 2018; Koricheva & Hayes, 2018), the extent to which genetic diversity can influence tree productivity through increased functional diversity is still unclear.

**Fig. 1.**
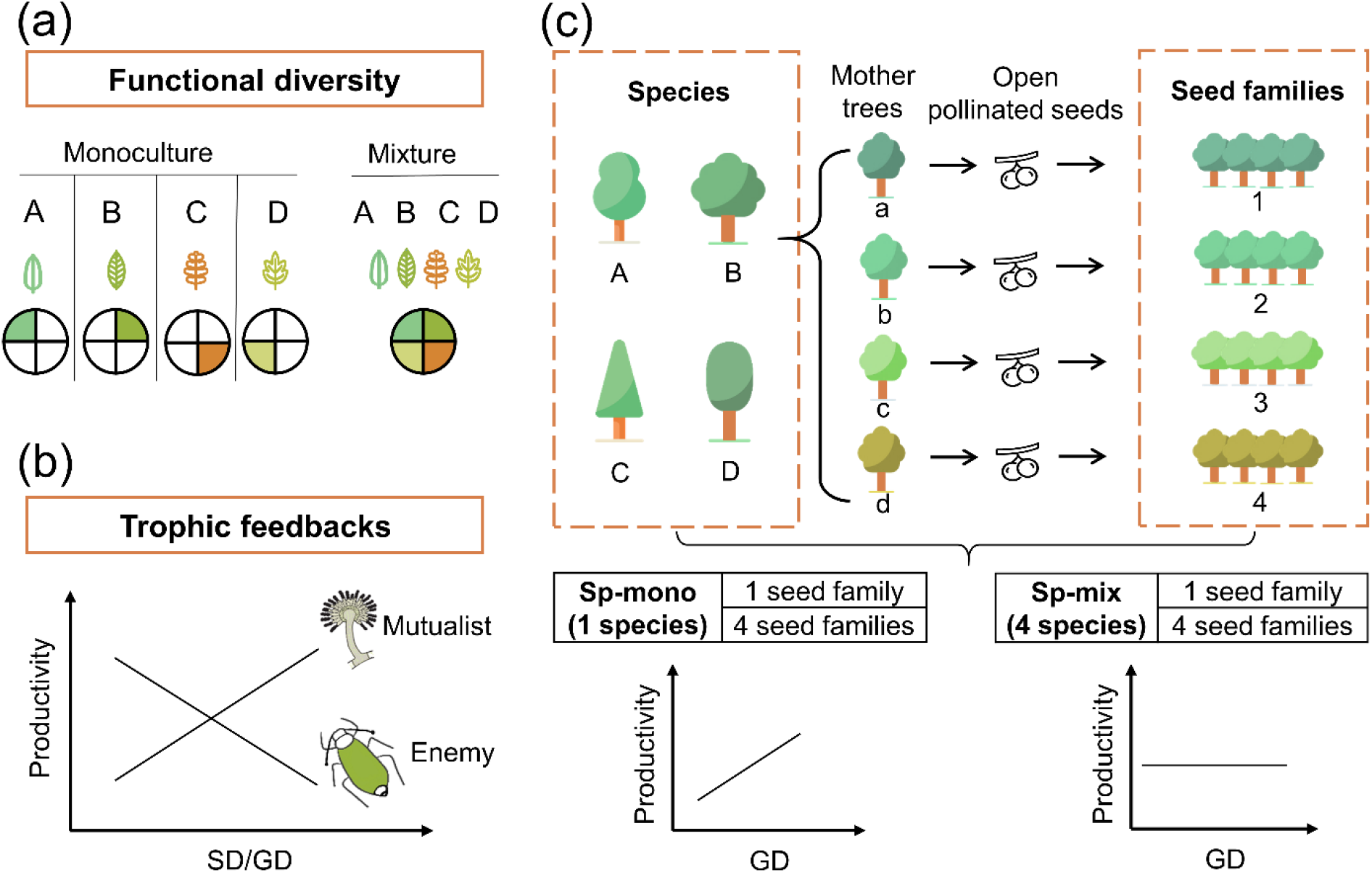
Conceptual illustration of the effects of functional diversity (a) and trophic feedbacks on tree productivity (b) and the experimental design (c). **(a)** shows resources for plant growth or other trophic groups in complementary ways due to functional diversity: the four hypothetical species/genotypes (A, B, C, D) with different functional traits (indicated by coloured leaves) are able to use a heterogeneous resource (indicated by coloured segments), thereby resulting in increased plant growth or providing niche opportunities for other trophic groups (Díaz & Cabido, 2001). **(b)** shows the mechanism of trophic feedbacks: with the increase of species diversity (SD) or genetic diversity (GD), negative feedbacks of enemies (e.g. herbivores) on tree productivity decrease due to diluted densities (Duffy, 2003) and positive feedbacks of mutualists on tree productivity increase due to increased diversity (e.g. mycorrhizal fungi (Semchenko et al., 2018)). **(c)** We represent tree species and genetic diversity by the number of species and seed families (all seeds from the same mother tree are defined as a single seed family), respectively. Species diversity and genetic diversity per plot were both 1 or 4, resulting in a full factorial design of species × genetic diversity. We hypothesize that the positive effects of tree genetic diversity should be stronger in tree species monocultures (Sp-mono) than mixtures (Sp-mix).

Trophic feedbacks, which result from the interactions of plants of different species/genotypes with other trophic groups, have been suggested as an additional mechanism underpinning positive biodiversity effects (Laforest-Lapointe et al., 2017). Trophic feedbacks can enhance the performance of a species mixture or a genotype mixture of a given species either by diluting the density of enemies (e.g. pathogens (Schmid, 1994) or herbivores (Jactel & Brockerhoff, 2007)) or enhancing diversity of beneficial mutualists (e.g. mycorrhizal fungi (Semchenko et al., 2018)) (Fig. 1b). These trophic feedbacks can be affected by plant functional diversity (Schuldt et al., 2019) and other factors (e.g. structural diversity (Schuldt et al., 2019)) which may provide more niche opportunities for other trophic groups. However, whereas many studies have analysed how plant diversity influences other trophic groups (Scherber et al., 2010; Schuldt et al., 2019) or how trophic interactions affect plant performance (Eisenhauer, 2012; Semchenko et al., 2018), effects of plant diversity on other trophic groups and the feedbacks of these on productivity have rarely been analyzed in combination.

In real world ecosystems, plant species diversity and genetic diversity can hardly be expected to influence ecosystems separately (Vellend & Geber, 2005). Previous studies of herbaceous plant communities have shown that the intensity of competition among species can be lowered by increased genetic diversity, which modifies the relationship between plant species diversity and plant productivity (Schöb et al., 2015). Likewise, the relative extent of plant intraspecific variation in functional traits, partly due to genetic diversity, has been shown to decrease with the increase of species diversity (Siefert et al., 2015). Although there are few forest experimental studies that manipulate both species and genetic diversity, most of them only compared their relative importance on ecosystem functions (Cook-Patton et al., 2011; Koricheva & Hayes, 2018), and we barely know their interactive effects via functional diversity and trophic feedbacks on plant productivity.

Here we disentangle how tree species diversity and genetic diversity affect tree community productivity via the impact of tree functional diversity and trophic feedbacks. We use data from a long-term tree species × genetic diversity experiment in a subtropical forest (Bruelheide et al., 2014) (BEF-China, www.bef-china.com). Tree species diversity (1 or 4 species per plot) and genetic diversity (1 or 4 seed families per species per plot) were manipulated in a factorial design to generate four plant diversity levels (Fig. 1c). We measured five morphological and chemical leaf traits, which have been shown to relate to resource acquisition (Cornelissen et al., 2003) and can have substantial variation both among and within species (Albert et al., 2010). Functional diversity was calculated as the variation of these five traits among seed families (Laliberté & Legendre, 2010). We quantified trophic interactions either by direct measurements of interactions (i.e. herbivory), or using the diversity of the trophic group (i.e. soil fungi) as a proxy to capture unspecific interactions potentially underpinning BEF relationships (Delgado-Baquerizo et al., 2016). Specifically, we tested whether tree species and genetic diversity increased tree community productivity via increased functional diversity (Fig. 1a) and trophic feedbacks (Fig. 1b). Furthermore, we tested whether effects of genetic diversity were more important in species monocultures than in species mixtures, because in the latter case genetic diversity between species may compensate for genetic diversity within species (Fig. 1c).

## Results

### Direct bivariate relationships between tree diversity, trophic interactions and tree community productivity

Overall, tree community productivity was significantly higher in 4-species mixture than in species monoculture (Fig. 2a), while genetic richness had no effect on tree productivity in the bivariate analyses (Fig. 2a). Tree functional diversity was higher in species mixture than species monoculture and was also higher in genetic mixture than genetic monoculture (Fig. 2b). The effects of genetic diversity on tree functional diversity, herbivore leaf damage and soil fungal diversity differed between species monocultures and species mixtures. Tree functional diversity in four seed-family species monocultures was larger than in one seed-family species monocultures but did not differ between species mixtures with four or one seed family per species (Fig. 2b). Additionally, both herbivore leaf damage and soil fungal diversity were similar in one and four seed-family species monocultures but lower in species mixtures with four than species mixtures with one seed family per species (Fig. 2c and d).

**Fig. 2.**
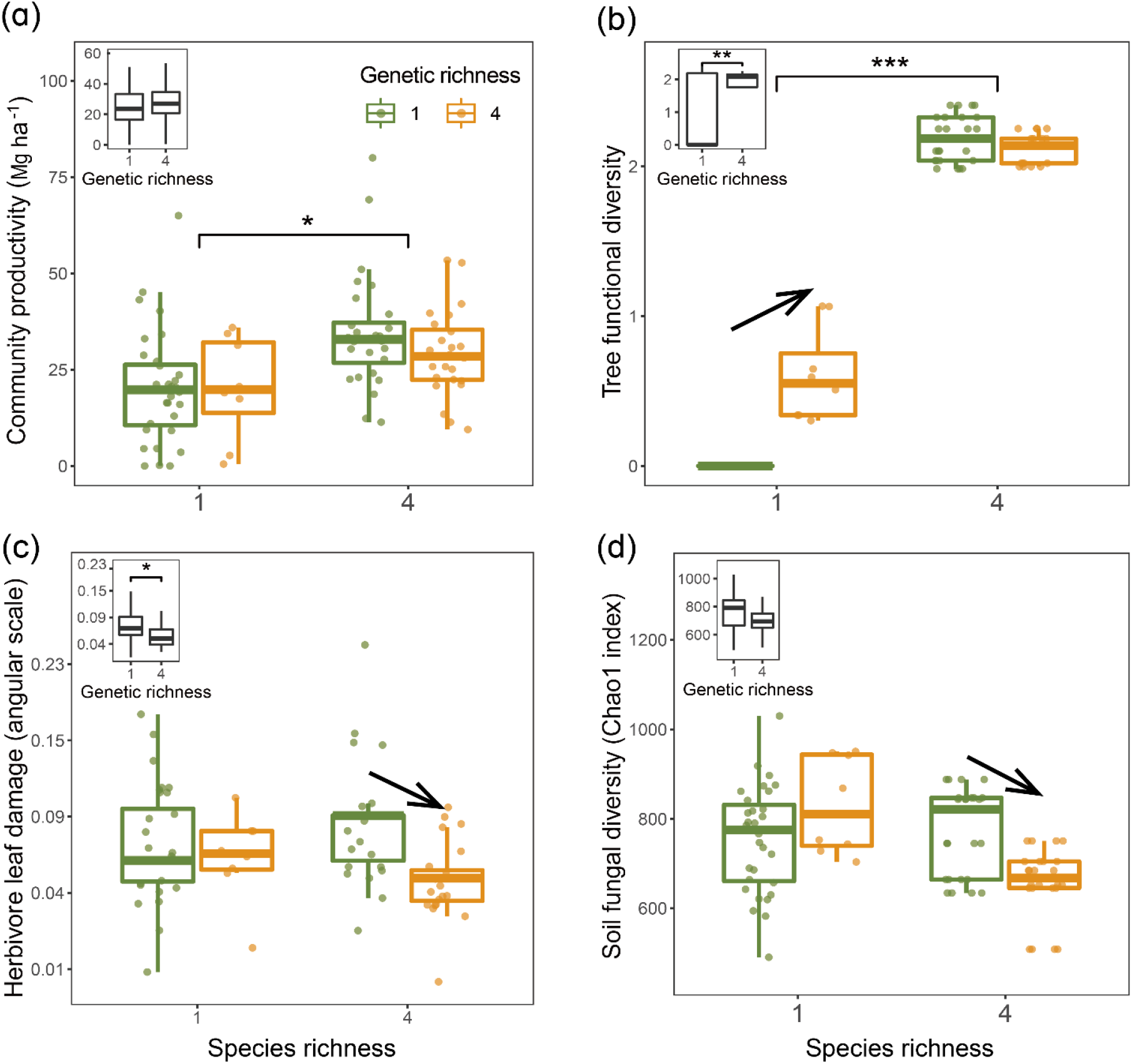
Tree community productivity, tree functional diversity and trophic interactions in tree communities of low vs. high species and genetic richness. The following effects were tested in LMMs: species richness main effect (left vs. right pair of bars in each panel), genetic richness main effect (inset on upper left in each panel), genetic richness effect within each species-richness level (arrows between pars within pairs). (a) tree community productivity, (b) tree functional diversity, (c) herbivore leaf damage, (d) soil fungal diversity. Asterisks indicate statistical significance (*** *P* < 0.0001, ** *P* < 0.001, * *P* < 0.05); solid arrow indicates *P* < 0.05,without arrow indicates *P* > 0.1). Details of the fitted models are shown in Table S4.

Tree functional diversity had a positive overall effect on community productivity but this effect was mainly due to an increase in functional diversity from species monocultures to mixtures (Fig 3a). Herbivore leaf damage and soil fungal diversity showed negative overall effects on tree productivity (marginally significant for herbivory and significant for fungal diversity; Fig. 3b, c). Furthermore, the effects of herbivore damage were different between genetic monocultures and genetic mixtures in species monoculture (Fig. 3b), while the effects of soil fungal diversity were different between genetic monocultures and genetic mixtures in species mixture (Fig. 3c).

**Fig. 3.**
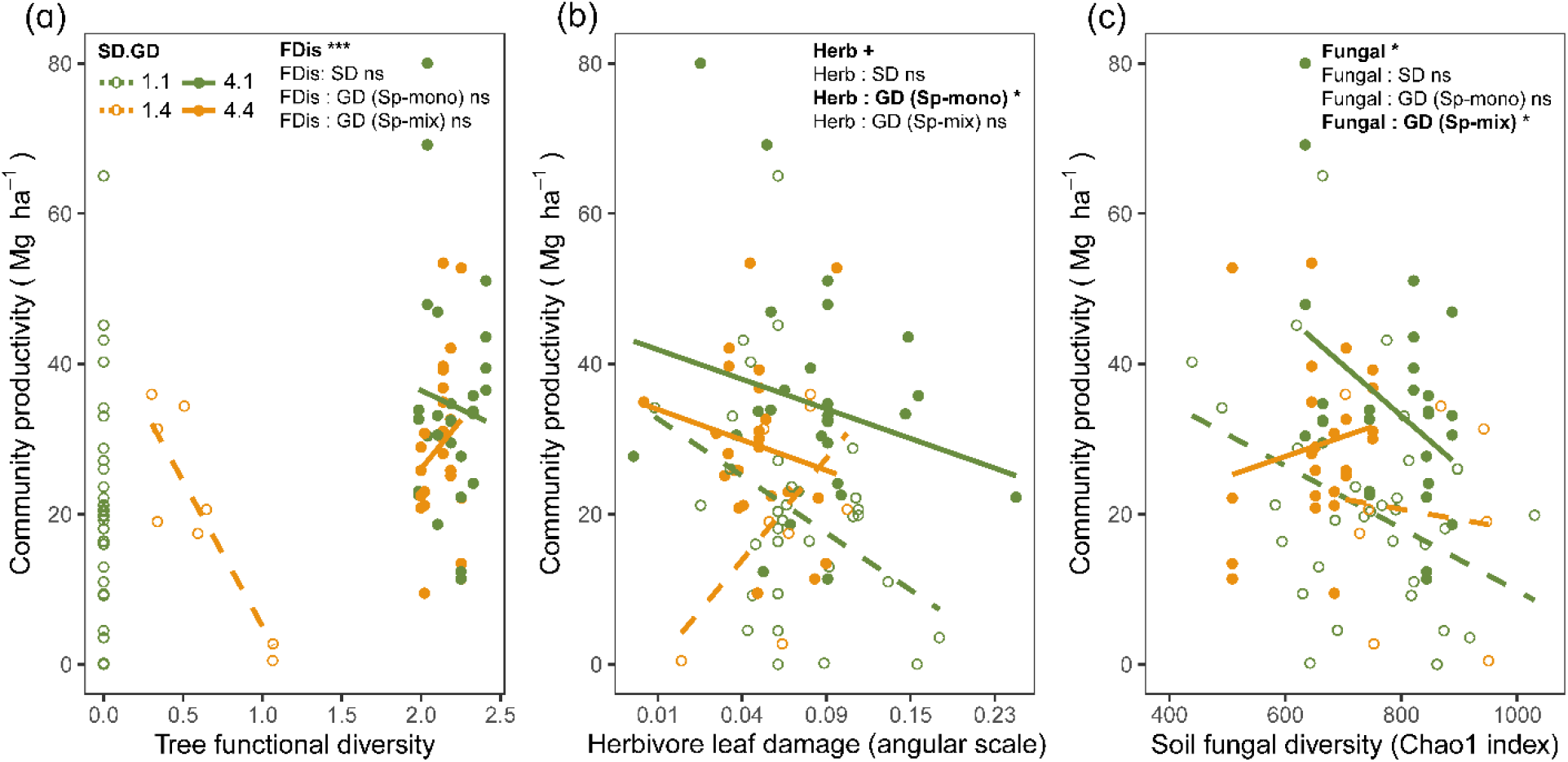
Bivariate relationships between tree community productivity and tree functional diversity (a), herbivory (b), and soil fungal diversity (c). Green dashed symbols represent genetic monocultures in species monoculture, green solid symbols represent genetic monocultures in species mixture, yellow dashed symbols represent genetic mixtures in species mixtures, orange solid symbols represent genetic mixture in species mixture. ‘FDis’ indicates tree functional diversity, ‘Herb’ indicates herbivore damage, ‘Fungal’ indicates soil fungal diversity, ‘Sp-mono’ presents species monocultures and ‘Sp-mix’ presents species mixtures, ‘SD’ indicates species diversity, ‘GD’ indicates genetic diversity and ‘:’ indicates the interaction effects. Asterisks indicate statistical significance (*** *P* < 0.0001, ** *P* < 0.001, * *P* < 0.05, + *P* < 0.1 and ns *P* > 0.1).

### Functional diversity and trophic feedbacks explain the effects of tree species and genetic diversity on tree productivity

Tree species and genetic diversity promoted tree community productivity as well as trophic interactions primarily indirectly through functional diversity (Fig. 4). The increase in functional diversity was larger for increasing species diversity than for increasing genetic diversity (standardized path coefficient 0.960 vs. 0.074, Fig. 4). Herbivory and soil fungal diversity reduced tree community productivity (Fig. 4, see also Fig. 3b, c). Overall, tree diversity had contrasting effects on tree community productivity through different mechanisms: tree species and genetic diversity promoted tree functional diversity which increased productivity directly but reduced it indirectly via negative feedbacks of herbivory and soil fungal diversity. However, species and genetic diversity also had positive indirect effects on community productivity via reduced soil fungal diversity (and genetic diversity additionally via reduced herbivory; Fig. 4). Whereas tree functional diversity and trophic feedbacks explain all effects of tree species diversity on productivity, there remained a direct negative effect of tree genetic diversity on productivity which could not be explained by the measured covariates (Fig. 4).

**Fig. 4.**
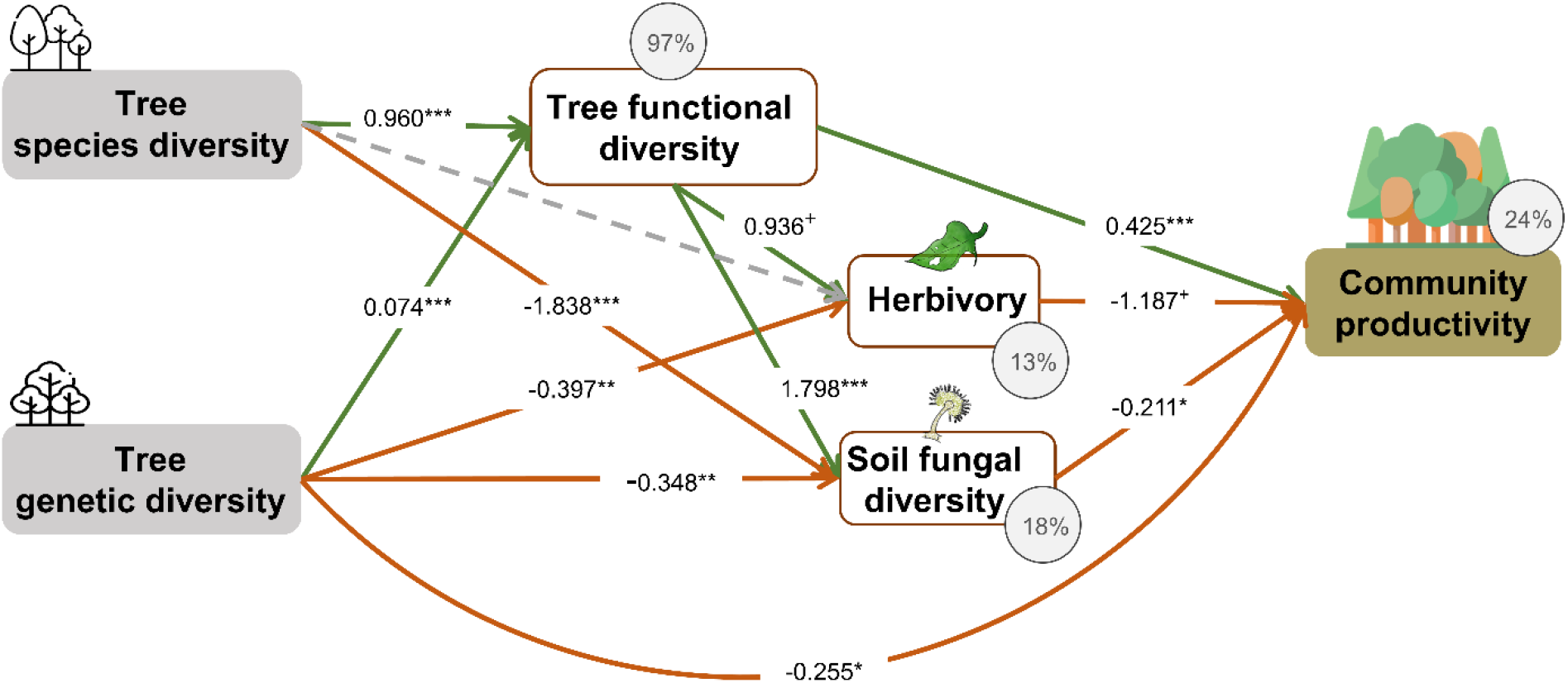
Effects of tree diversity on higher trophic levels and tree community productivity (global Fisher’s C = 1.677, DF = 4, *P* = 0.795, initial AICc = 56.840, final AICc = 54.004). Positive and negative paths are indicated in green and orange. The standardized path coefficients are given by the numbers, statistical significance is indicated by asterisks (*** *P* < 0.0001, ** *P* < 0.001, * *P* < 0.05 and + *P* < 0.1) and the explained variance of dependent variables is indicated by the percentage values. Grey dashed lines indicate non-significant (*P* > 0.1) pathways in the final model.

### Effects of tree genetic diversity in species monocultures and species mixtures

When the above analysis was split into two (Fig. 5), we found that tree genetic diversity negatively affected community productivity via functional diversity in species monocultures and had no such effect in species mixtures (see also Fig. 2b and Fig. 3a). Positive indirect effects through herbivory (resulting from two negative paths from genetic diversity to herbivory and form herbivory to community productivity) were similar in both species monocultures and mixtures. Tree genetic diversity increased soil fungal diversity and tree productivity in species monocultures but reduced them in species mixtures, resulting in negative indirect effects on community productivity in the first (Fig. 5a) and positive indirect effects on community productivity in the second case (Fig. 5b). The negative indirect effect of genetic diversity on community productivity via functional diversity in species monocultures, which contrasts with the combined analysis, was counterbalanced by a positive direct effect of genetic diversity on productivity, indicating that here other aspects than those included with the five functional traits measured were important.

**Fig. 5.**
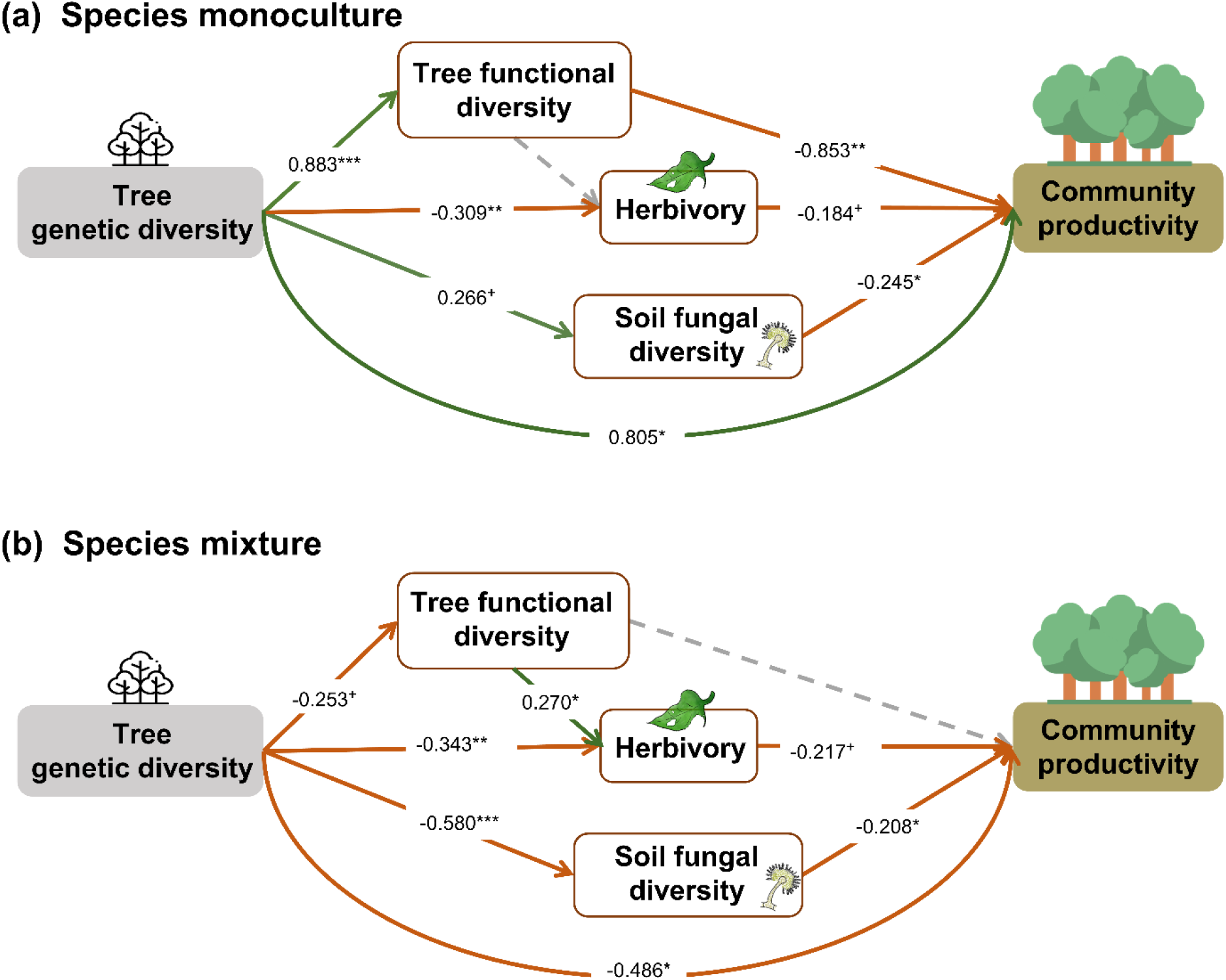
Effects of tree genetic diversity on higher trophic levels and tree community productivity in tree species monocultures (a) and mixtures of four tree species (b). global Fisher’s C = 3.416, DF = 4, *P* = 0.491, initial AICc = 45.704, final AICc = 43.528. Positive and negative paths are indicated in green and orange. The standardized path coefficients are given by the numbers, and statistical significance is indicated by asterisks (*** *P* < 0.0001, ** *P* < 0.001, * *P* < 0.05 and + *P* < 0.1). Grey dashed lines indicate non-significant (*P* > 0.1) pathways in the final model.

## Discussion

Our study showed that manipulating tree species and genetic diversity in a factorial design reveals strong effects of both as well as their interaction on measured ecosystem variables. Nevertheless, effects of genetic diversity on tree community productivity through functional diversity were weaker than those of genetic diversity on trophic interactions (see Fig. 4), indicating that mechanisms underpinning effects of genetic diversity may in part differ from those underpinning effects of species diversity, as we will discuss below. Regarding our two hypotheses, we found that tree species and genetic diversity can increase tree community productivity via increased functional diversity and trophic feedbacks as predicted by our first hypothesis. This suggests complementary resource-use and biotic niches, respectively, as mechanisms underpinning the biodiversity effects (Turnbull et al., 2016). Furthermore, as predicted by our second hypothesis, we found that the effects of tree genetic diversity on productivity were different in tree species monocultures than in mixtures. In the following, we discuss these results in more detail.

### Tree species and genetic diversity drive tree community productivity via functional diversity and trophic feedbacks

The observed tree biodiversity effects via increased functional diversity on tree community productivity were both positive and direct, and negative and indirect via changed trophic feedbacks (see Fig. 4). The positive direct effect indicates that tree functional diversity enhanced complementary resource acquisition at tree community level (Kahmen et al., 2006; Marquard et al., 2009; Williams et al., 2017), which consequently enhanced tree community productivity. Whereas negative effects of herbivory on plant productivity via the reduction of leaf area (Zvereva et al., 2012) and photosynthesis of remaining leaves (Nabity et al., 2009) are to be expected. The negative effects of soil fungal diversity on productivity correspond with the finding that the majority of these fungi were saprophytes (Fig. S1), competing with plants for resources (Kaye & Hart, 1997; van der Heijden et al., 2008). Indeed, in a related study in the same region, the diversity of saprophytic fungi has been found to decrease multifunctionality (Schuldt et al., 2018).

Meanwhile, tree functional diversity also provided more niche opportunities to benefit generalist herbivores and soil fungi which reduced tree community productivity, as has been found for these tree species in a parallel field study nearby (Brezzi et al., 2017). Indirect positive effects of species and genetic diversity—remaining after paths via functional diversity—via reduced herbivory and soil fungal diversity further increased community productivity (see Fig.4). This finding corresponds to previous studies that found that plant diversity could reduce negative feedbacks of other trophic groups by decreasing the density and diversity of specialist enemies (e.g. Duffy, 2003; Jactel & Brockerhoff, 2007).

Even after accounting for tree functional diversity and trophic feedbacks, we still detected a direct negative effect of tree genetic diversity on tree productivity, while the direct effect of tree species diversity was fully explained by functional diversity and trophic feedbacks. This suggests that aspects of genetic diversity that do not contribute to functional diversity or trophic interactions as measured in this study may reduce ecosystem functioning, e.g. due to trade-offs between genetic diversity and species diversity. For example, it has been shown that in species-diverse grassland ecosystems niche-complementarity between species can increase at the expense of reduced variation within species (van Moorsel et al., 2018; van Moorsel et al., 2019; Zuppinger-Dingley et al., 2014; Zvereva et al., 2012). Thus, our experiment simulating high genetic diversity within species in mixtures might have reduced the positive effects of high species diversity. This interpretation would be compatible with the observation that in the separate path analyses direct negative effects of genetic diversity on productivity were only found in species mixtures, whereas in the species monocultures these effects were positive (see next section). Independent of this interpretation, our finding could also imply that partly different mechanisms underpin effects of species vs. genetic diversity on ecosystem functioning (Barantal et al., 2019; Des Roches et al., 2018).

### Effects of tree genetic diversity differ between tree species monocultures and mixtures

As expected, the effects of tree genetic diversity via functional diversity on tree community productivity were stronger in tree species monocultures than mixtures. However, these effects in monocultures were negative rather than positive and were counterbalanced by positive direct effects of tree genetic diversity on productivity (see Fig. 5). Thus, other aspects of tree genetic diversity seem to play an important role not only for productivity in tree species mixtures (see previous section) but also for productivity in tree species monocultures. These may include unmeasured functional traits such as root traits (Bardgett et al., 2014) or unknown mechanisms underpinning effects of tree genetic diversity. Effects of tree genetic diversity on soil fungal diversity were of similar strength but again partly opposing direction in tree species monocultures vs. mixtures (see Fig. 2 and Fig. 5). While it is difficult to speculate about potential causes of these differences in tree genetic diversity effects between tree species mixtures and monocultures, they again hint at trade-offs between genetic and species diversity as discussed in the previous section.

## Conclusion

This study tried to disentangle effects of tree species and genetic diversity via functional diversity and trophic feedbacks on tree community productivity in a simple experimental system with 4 species and multiple seed families per species. Even though this was already challenging to set up, manage and assess by measurements on trees and soil samples, larger studies will be required to generalize results. Nevertheless, our results suggest that both partitioning of resource-use and enemy niches (Turnbull et al., 2016) between and among genotypes within tree species played a role in affecting tree community productivity. Although both tree species and genetic diversity contributed to productivity, the underpinning mechanisms differed and were harder to explain for tree genetic diversity. We suggest that trade-offs between tree species and genetic diversity may cause the latter to switch strength and direction between species monocultures and mixtures. Thus, while genetic diversity (within species) may be beneficial in monocultures and low-species mixtures it may have opposite effects in richer species mixtures. We were not able to definitively report causality between trophic feedbacks and tree productivity because we did not experimentally manipulate herbivore leaf damage and soil fungi. However, our results do support the hypothesis that trophic feedbacks affect plant community productivity. Given the importance of afforestation projects to mitigate carbon loss and provide ecological and economic benefits (Brockerhoff et al., 2008; Lamb et al., 2005), we strongly recommend that both tree species and genetic diversity should be considered in afforestation projects.

## Materials and Methods

### Study site and experimental design

This study was carried out in the species × genetic diversity experiment of the Biodiversity– Ecosystem Functioning Experiment China Platform (BEF-China, www.bef-china.com; Bruelheide et al., 2014; Hahn et al., 2017). BEF-China is located close to Xingangshan, Dexing City, Jiangxi Province, China. The mean annual temperature is 16.7 °C and mean annual precipitation is 1821 mm. The species × genetic diversity experiment was established in 2010 and comprises 24 plots of 25.8 × 25.8 m^2^ equal to one Chinese unit of “mu”). Each plot was planted with 400 individual trees from a pool of four species (*Alniphyllum fortunei*, *Cinnamanum camphora*, *Daphniphyllum oldhamii* and *Idesia polycarpa*) with the mother trees of all tree individuals known. We defined the offspring from the same mother tree as a seed family and we assumed that the genetic variation is larger among seed families than within a seed family (Bongers et al., 2020; Hahn et al., 2017). Since the offspring of a single mother tree could have been sired by different father trees, they represented anything between full- and half-sib families. Thus, in this study, we used the number of seed families per species as a measure of genetic diversity (Bruelheide et al., 2014). Across the 24 plots, we combined species diversity (1 or 4 species) and genetic diversity (1 or 4 seed families per species) which resulted in four tree diversity levels: one species with one seed family (1.1), one species with four seed families (1.4), four species with one seed family per species (4.1) and four species with four seed families per species (4.4) (Bongers et al., 2020). The 1.1 and 1.4 diversity treatments were applied at subplot level (0.25 mu) and replicated 8 and 2 times per species, respectively. The 4.1 and 4.4 diversity treatments were applied at plot level (1 mu) and were replicated 8 and 6 times, respectively (Bongers et al., 2020; Bruelheide et al., 2014). To allow for simpler analysis, we obtained most community measures at subplot level also for the 4.1 and 4.4 diversity treatments and thereafter used the subplots for all tests of diversity effects on these community measures. In total, because one 1-mu plot could not be established due to logistic constraints, the number of subplots used was 92 (32 subplots of 1.1, 8 subplots of 1.4, 28 subplots of 4.1 and 24 subplots of 4.4 diversity treatment).

### Tree functional traits and functional diversity

Five leaf functional traits were measured in 2017 and 2018, including leaf area (LA), specific leaf area (SLA), chlorophyll content (CHL), leaf nitrogen content (LN) and leaf carbon content (LC). These traits can reflect the resource acquisition ability of plants and may show substantial variation not only among species but also within species (Albert et al., 2010; Cornelissen et al., 2003). We collected these traits on 547 individuals across all the species × genetic diversity combinations (Table S1), with details described in Bongers et al. (2020).

Functional leaf trait diversity was expressed as multivariate functional dispersion (FDis) (Laliberté & Legendre, 2010), which we used to quantify the mean distance of individual seed families to the centroid of all seed families in the community (Laliberté & Legendre, 2010). In every mixture trait values were weighted equally across seed families and species because these were planted in equal numbers in each plot or subplot. The mean value of FDis per species × genetic diversity level was used to fill in missing values in a few subplots with families lacking trait data (Table S1). We also calculated another frequently used functional diversity index, Rao’s Q (Rao, 1982). However, a strong positive correlation was detected between FDis and Rao’s Q in simulated data (Laliberté & Legendre, 2010) and in our study (Fig. S2). Moreover, in the case of equal weighting, FDis should perform better than Rao’s Q (Laliberté & Legendre, 2010). Therefore, we only used FDis in the analyses presented in this study. The calculations of FDis and Rao’s Q were done with the ‘dbFD’ function of the ‘FD’ R package (Laliberté et al., 2014).

### Trophic interactions

#### Herbivory

Herbivory results from the interaction between plants and herbivores and can be recorded as leaf damage. For every individual tree, four or five damaged leaves were randomly collected and herbivory visually estimated (Johnson et al., 2016) (same 547 trees as for the traits, see above) in 2017. Thus, in this study herbivory represents the percentage of damaged area per leaf attacked by herbivores. The herbivory caused by chewers, gall formers, leaf miners and rollers were collectively counted. Because we only collected damaged leaves in this study, we might have overestimated the herbivory per individual tree. We therefore used data from other plots of the BEF-China experiment (Schuldt et al., 2015) to correct the potential bias. We related leaf damage including non-damaged leaves (total leaf damage) to leaf damage excluding non-damaged leaves (damage per damaged leaf) for all four species by linear regression (Pearson’s correlation = 0.86– 0.96, *P* < 0.001) (Table S2). With these regression models, we got the predicted values of herbivory for our observational data and used these predicted values in the final analyses. The mean value of herbivore damage per species × genetic diversity level was used to fill in missing values in a few subplots with tree individuals lacking herbivory data (Table S1).

#### Soil fungal diversity

Soil fungal diversity was used as proxies of unspecified trophic interactions. Soil samples were taken on subplot level for the 1.1 and 1.4 diversity treatments, but on plot level for the 4.1 and 4.4 diversity treatments in 2017. In each subplot or plot, five soil samples from the top 0–5 cm soil layer were collected from the four corners and the center of each subplot or plot. The five samples were then mixed together. Each soil sample was packed with dry ice and transferred to the laboratory for storage at −80°C until DNA extraction. The total genomic DNA of the subsample was extracted using Soil Genomic DNA Kit (Tiangen Biotech Co., Beijing, China), following the manufacturer’s protocol. The DNA was extracted to perform PCR amplification. We amplified the nuclear rDNA internal transcribed spacer 2 (ITS2) region using primers ITS3F (GCATCGATGAAGAACGCAGC) and ITS4R (TCCTCCGCTTATTGATATGC). We processed the raw sequences with the QIIME 2 pipeline (https://docs.qiime2.org/) to cluster and assign operational taxonomic units (OTU). The fungal OTU tables were rarefied to 10975 reads to account for the different sequencing depths. We then assigned the sequences to taxonomic groups using the UNITE database (Nilsson et al., 2019). Based on the taxonomic and abundance information of every subplot or plot, the Chao1 diversity index (Chao, 1984) was used to quantify soil fungal diversity, because most fungal species in our study were relatively rare and the Chao1 index can account well for rare species (Chao, 1984). The calculation of diversity of soil fungi was done with the ‘vegan’ package version 2.5-7 in R (Oksanen et al., 2019).

#### Tree community productivity

We measured the basal area (BA) and the height (H) of all trees in the species × genetic diversity plots in 2018 (Bongers et al., 2020). Individual tree biomass (kg) was calculated using the biomass equation (H × BA × CV) of the BEF-China experiment (Huang et al., 2018) in which CV is a correction factor for stem shape and wood density. More detail about the biomass equation can be found in Huang et al. 2018. We summed the biomass of individual trees to subplot level to calculate tree community productivity (Mg ha^-1^).

### Statistical analysis

First, we evaluated the bivariate relationships between tree diversity, trophic interactions and tree community productivity. To determine how species and genetic diversity and their interaction affected tree functional diversity and trophic interactions, linear mixed-effects models (LMMs) were fitted with the sequential order of the fixed-effects explanatory terms species diversity, genetic diversity and the interaction of them. Additionally, to detect the effects of genetic diversity in species monocultures and species mixtures separately, we also fitted the model by contrast codes (Table S3) to indicate the genetic diversity only with species monocultures or species mixtures. For all the linear mixed-effects models, we used ‘plot’ as a random variable since subplots were nested in plots. LMMs were fitted with the ‘lmer’ function of the lme4 package version 1.1.27.1 (Bates et al., 2015) and Kenward-Roger’s method was used to calculate denominator degrees of freedom and F-statistics by lmerTest-package version 3.1.3 (Kuznetsova et al., 2017). To meet the assumptions of linear mixed models, the proportion of leaf damage caused by herbivores was angular transformed (George W. Snecdecor, 1989). We also tested the effects of tree functional diversity, herbivore leaf damage and soil fungal diversity, and the interaction effects of them with species diversity and genetic diversity in both species monocultures and species mixtures on tree community productivity using linear models with the functions ‘lm’ in R (www.r-project.org).

Second, we fitted path models (Grace, 2006) with the ‘piecewiseSEM’ package version 2.1.2 in R (Lefcheck, 2016) to assess causal hypotheses about how the effects of tree species and genetic diversity on community productivity could have been mediated via tree functional diversity and trophic interactions. The initial model was constructed by the most relevant pathways derived from theoretical assumptions (Fig. S3). Additionally, we used separate linear regressions to assess relationships between variables hypothesized to be related in cause–effect relationships in the path models. We assumed that both tree genetic diversity and species diversity could influence trophic interactions and community productivity directly or indirectly, i.e., mediated via functional diversity (Müller et al., 2018; Scherber et al., 2010; Schuldt et al., 2019). Moreover, we hypothesized that tree functional diversity, herbivore leaf damage and soil fungal diversity have direct feedbacks on community productivity (Eisenhauer, 2012; Semchenko et al., 2018). We sequentially dropped non-informative pathways, if their removal reduced the AICc of the path models (Grace, 2006).

Thirdly, separate multi-group path models were fitted for species monocultures and mixtures, since significant interactions between species and genetic diversity in the ANOVAs indicated that genetic diversity had different effects in species monocultures vs. mixtures. The initial multigroup model is shown in Fig. S4. We simplified the multigroup initial model with the same procedure as in the above second part by comparing AICc values (Grace, 2006).

All the analyses were carried out in R 4.0.5 (www.r-project.org).

## Acknowledgements

We acknowledge the support of the BEF-China platform. This study was financially supported by the Strategic Priority Research Program of the Chinese Academy of Sciences (XDB31000000), the National Natural Science Foundation of China (31870409 and 32161123003). X.L. was supported by the Youth Innovation Promotion Association CAS (2019082). B.S. was supported by the University Research Priority Program Global Change and Biodiversity of the University of Zurich.

## Data availability

All numerical data were used to generate the figures that have been deposited in Dryad. During the review period, the dataset could be downloaded by this temporary link: https://datadryad.org/stash/share/WOhia74fAe8WuUcqk0Tvxw6N91SOlyQmjaQwJ6fpWyk.

## Supplementary files

The following Supporting Information is available for this article:

---

**Fig. S1.**
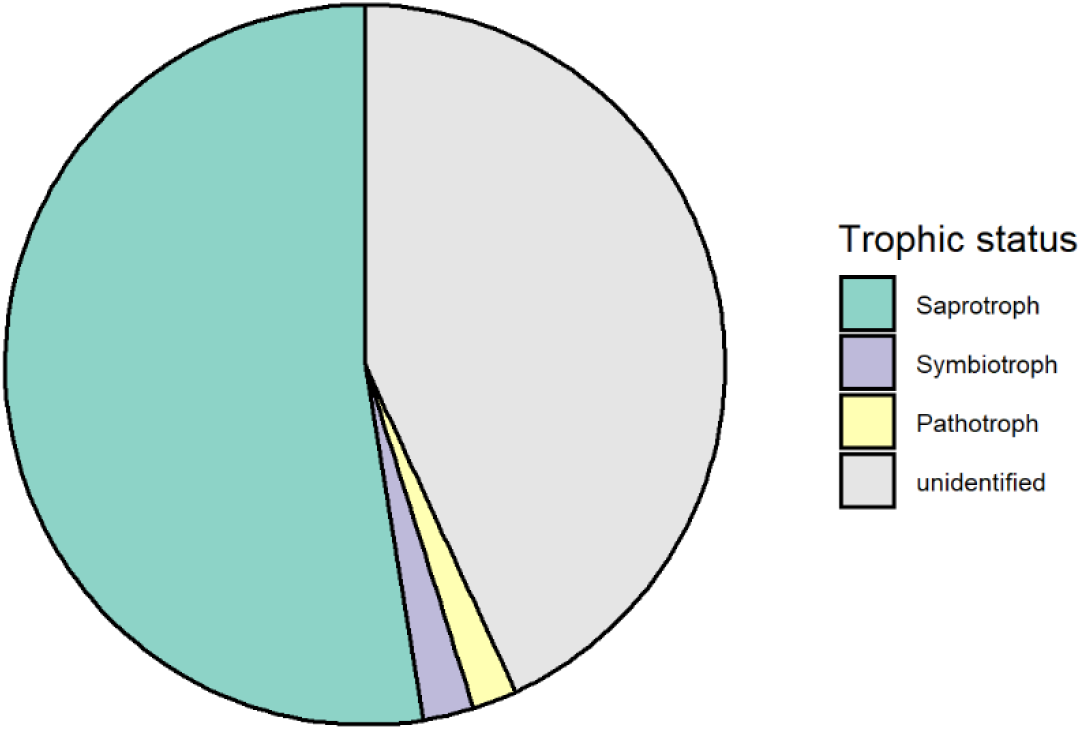
Trophic composition of soil fungi in this study. All fungi from this study were pooled together to calculate the relative abundance of each trophic group.

**Fig. S2.**
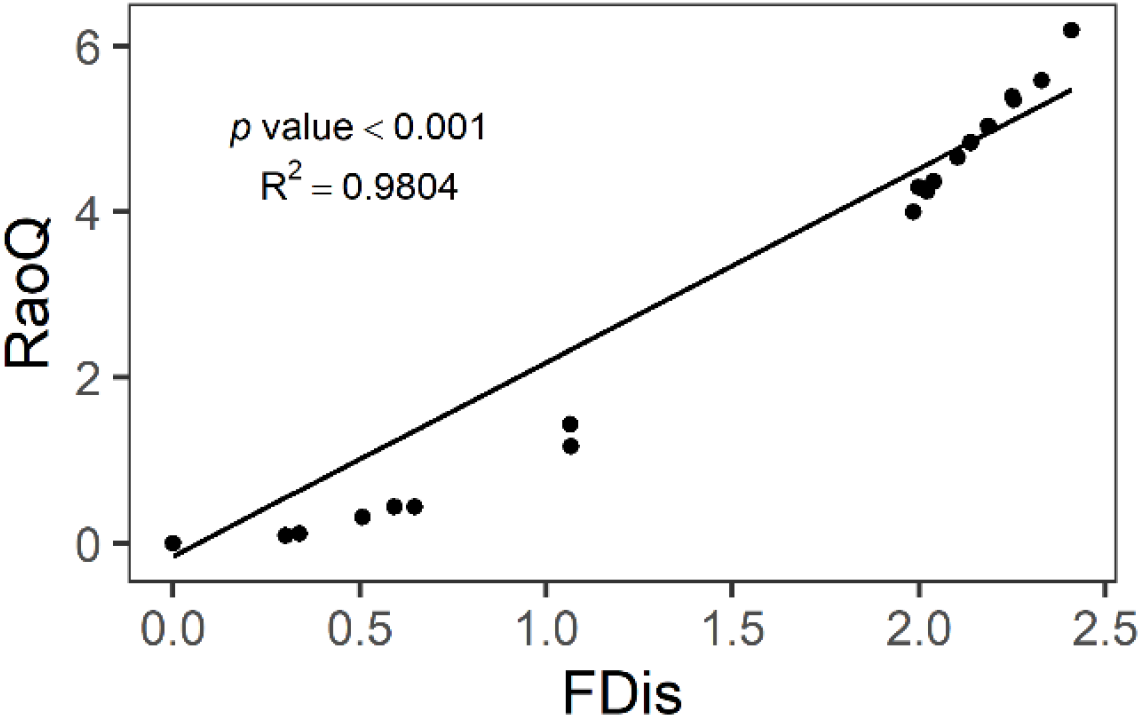
Relationship between functional dispersion (FDis) and Rao’s Q (RaoQ).

**Fig. S3.**
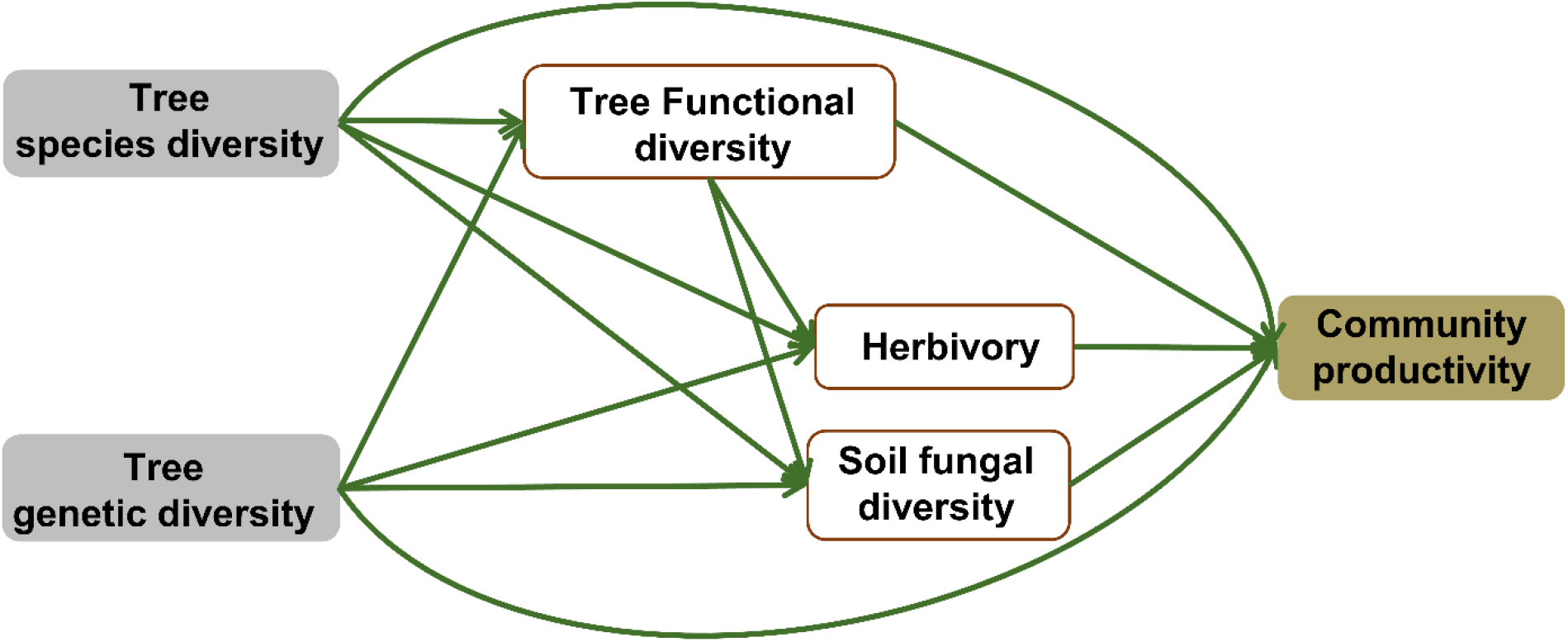
Initial path model used in the present study. Tree species diversity represents the number of tree species, genetic diversity represents the number of seed families per tree species, functional diversity (based on FDis) represents the mean distance of individual species seed families to the centroid of all species seed families in a subplot. Herbivory represents the percentage of herbivore leaf damage, soil fungal diversity was quantified by Chao1 diversity index, community productivity represents tree productivity of subplots (Mg ha^-1^).

**Fig. S4.**
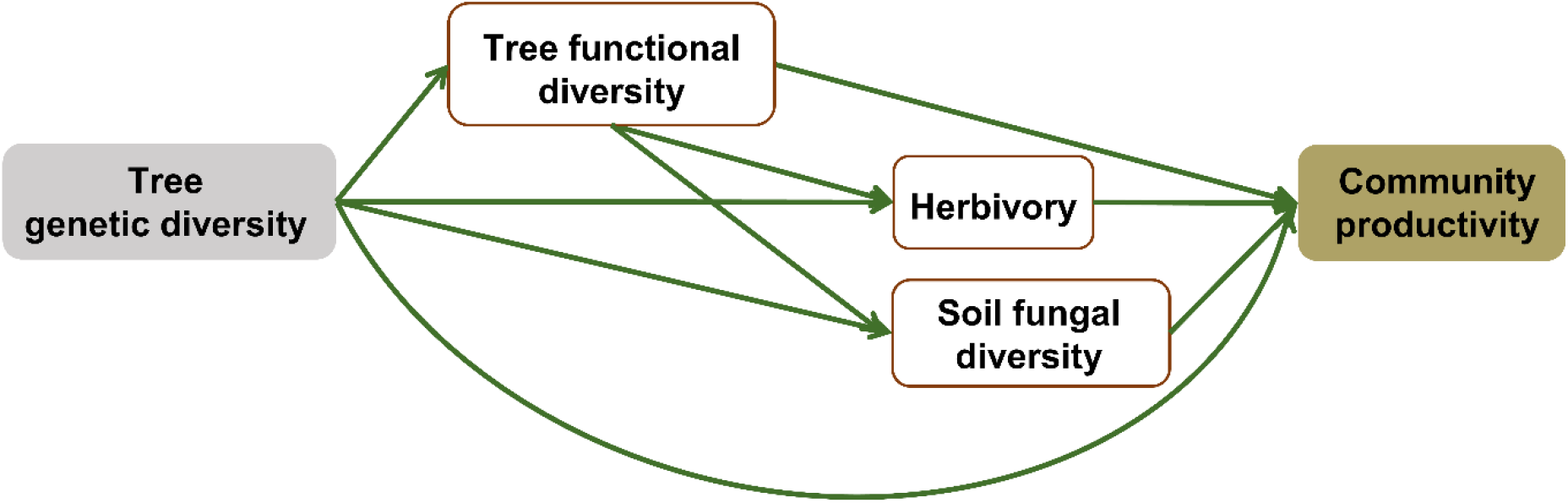
Initial path model structure of genetic diversity effects in both species monocultures and mixtures. Genetic diversity represents the number of seed families per tree species. Functional diversity (based on FDis) represents the mean distance of individual species seed families to the centroid of all species seed families in a subplot. Herbivory represents the percentage of herbivore leaf damage, soil fungal diversity was quantified by Chao1 diversity index, community productivity represents tree productivity of subplots (Mg ha^-1^).

**Table S1.** Data description of multi-trophic levels.

| Data type | Data description | Subplots | Year |
| --- | --- | --- | --- |
| Plant trait | LA, SLA, CHL, LN, LC | 77 | 2017 |
| Herbivore damage | Visually estimated | 77 | 2017 |
| Soil fungi | Mainly composed of saprophytes | 53 | 2017 |
| Community productivity | Sum of the biomass per subplot/area of subplot | 92 | 2018 |

**Table S2.** Results of linear models of leaf damage excluding undamaged leaves (this study) ∼ leaf damage including undamaged leaves (from other plots of the BEF-China experiment) for the four species used in the present study. These models were used to correct the potential bias of herbivory estimates as a result of only collecting damaged leaves.

| Species | Slope | Intercept | R<sup>2</sup> | Pearson's correlation |
| --- | --- | --- | --- | --- |
| *Alniphyllum fortunei* | 0.89970 | 2.83483 | 0.86 | 0.93 |
| *Cinnamanum camphora* | 0.94465 | 2.07484 | 0.73 | 0.86 |
| *Daphniphyllum oldhamii* | 0.88387 | 2.54032 | 0.92 | 0.96 |
| *Idesia polycarpa* | 0.92406 | 1.77523 | 0.86 | 0.93 |

**Table S3.** Contrast coding of genetic diversity in species monocultures and species mixtures separately. Sp-mono presents species monocultures and Sp-mix presents species mixtures.

| Speices diversity | Genetic diversity | Genetic diversity in Sp-mono | Genetic diversity in Sp-mix |
| --- | --- | --- | --- |
| 1 | 1 | 1 | 0 |
| 4 | 1 | 0 | 1 |
| 1 | 4 | 4 | 0 |
| 4 | 4 | 0 | 4 |

**Table S4.** Summary of linear mixed-effects models (LMMs) of species diversity (SD), genetic diversity (GD), and their interactions on tree productivity, tree functional diversity and trophic interactions; and expressed values are Df representing degree of freedom and F values with related significances, ‘***’ indicates P < 0.001; ‘**’ indicates P < 0.01; ‘*’ indicates P < 0.05.

| Factors | Df | Tree community<br>productivity<br>F value | Tree functional<br>diversity<br>F value | Herbivory<br>F value | Soil fungal<br>diversity<br>F value |
| --- | --- | --- | --- | --- | --- |
| SD | 1 | **6.16\*** | **2137.67\*\*\*** | 0.08 | 1.06 |
| GD | 1 | 0.51 | **11.14\*\*** | **5.86\*** | 1.57 |
| SD×GD | 1 | 0.23 | **44.03\*\*\*** | 1.44 | **5.29\*** |
| SD | 1 | **6.16\*** | **2137.67\*\*\*** | 0.08 | 1.06 |
| GD (Sp-mono) | 1 | 0.00 | **53.99\*\*\*** | 0.17 | 1.35 |
| GD (Sp-mix) | 1 | 0.74 | 1.19 | **7.13\*** | **5.51\*** |

## References

1. Albert, C. H., Thuiller, W., Yoccoz, N. G., Douzet, R., Aubert, S., & Lavorel, S. (2010). A multi-trait approach reveals the structure and the relative importance of intra- vs. interspecific variability in plant traits. Functional Ecology, 24(6), 1192–1201. https://doi.org/10.1111/j.1365-2435.2010.01727.x

2. Barantal, S., Castagneyrol, B., Durka, W., Iason, G., Morath, S., & Koricheva, J. (2019). Contrasting effects of tree species and genetic diversity on the leaf-miner communities associated with silver birch. Oecologia, 189(3), 687–697. https://doi.org/10.1007/s00442-019-04351-x

3. Bardgett, R. D., Mommer, L., & De Vries, F. T. (2014). Going underground: root traits as drivers of ecosystem processes. Trends in Ecology & Evolution, 29(12), 692–699. https://doi.org/10.1016/j.tree.2014.10.006

4. Bates, D., Mächler, M., Bolker, B., & Walker, S. (2015). Fitting Linear Mixed-Effects Models Using lme4. Journal of Statistical Software, 67(1), 1–48. https://doi.org/10.18637/jss.v067.i01

5. Bongers, F. J., Schmid, B., Durka, W., Li, S., Bruelheide, H., Hahn, C. Z., Yan, H., Ma, K., & Liu, X. (2020). Genetic richness affects trait variation but not community productivity in a tree diversity experiment. New Phytologist, 227(3), 744–756. https://doi.org/doi:10.1111/nph.16567

6. Brezzi, M., Schmid, B., Niklaus, P. A., & Schuldt, A. (2017). Tree diversity increases levels of herbivore damage in a subtropical forest canopy: evidence for dietary mixing by arthropods? Journal of Plant Ecology, 10(1), 13–27. https://doi.org/10.1093/jpe/rtw038

7. Brockerhoff, E. G., Jactel, H., Parrotta, J. A., Quine, C. P., & Sayer, J. (2008). Plantation forests and biodiversity: oxymoron or opportunity? Biodiversity and Conservation, 17(5), 925–951. https://doi.org/10.1007/s10531-008-9380-x

8. Bruelheide, H., Nadrowski, K., Assmann, T., Bauhus, J., Both, S., Buscot, F., Chen, X. Y., Ding, B. Y., Durka, W., Erfmeier, A., Gutknecht, J. L. M., Guo, D. L., Guo, L. D., Hardtle, W., He, J. S., Klein, A. M., Kuhn, P., Liang, Y., Liu, X. J., Michalski, S., Niklaus, P. A., Pei, K. Q., Scherer-Lorenzen, M., Scholten, T., Schuldt, A., Seidler, G., Trogisch, S., von Oheimb, G., Welk, E., Wirth, C., Wubet, T., Yang, X. F., Yu, M. J., Zhang, S. R., Zhou, H. Z., Fischer, M., Ma, K. P., & Schmid, B. (2014). Designing forest biodiversity experiments: general considerations illustrated by a new large experiment in subtropical China. Methods in Ecology and Evolution, 5(1), 74–89. https://doi.org/Doi 10.1111/2041-210x.12126

9. Cadotte, M. W., Carscadden, K., & Mirotchnick, N. (2011). Beyond species: functional diversity and the maintenance of ecological processes and services. Journal of Applied Ecology, 48(5), 1079–1087. https://doi.org/10.1111/j.1365-2664.2011.02048.x

10. Cardinale, B. J., Duffy, J. E., Gonzalez, A., Hooper, D. U., Perrings, C., Venail, P., Narwani, A., Mace, G. M., Tilman, D., Wardle, D. A., Kinzig, A. P., Daily, G. C., Loreau, M., Grace, J. B., Larigauderie, A., Srivastava, D. S., & Naeem, S. (2012). Biodiversity loss and its impact on humanity. Nature, 486(7401), 59–67. https://doi.org/10.1038/nature11148

11. Ceballos, G., Ehrlich, P. R., Barnosky, A. D., Garcia, A., Pringle, R. M., & Palmer, T. M. (2015). Accelerated modern human-induced species losses: Entering the sixth mass extinction. Science Advances, 1(5), e1400253. https://doi.org/10.1126/sciadv.1400253

12. Chao, A. (1984). Non-parametric estimation of the number of classes in a population. Scandinavian Journal of Statistics, 11, 265–270

13. Cook-Patton, S. C., McArt, S. H., Parachnowitsch, A. L., Thaler, J. S., & Agrawal, A. A. (2011). A direct comparison of the consequences of plant genotypic and species diversity on communities and ecosystem function. Ecology, 92(4), 915–923. https://doi.org/10.1890/10-0999.1

14. Cornelissen, J. H. C., Lavorel, S., Garnier, E., Díaz, S., Buchmann, N., Gurvich, D. E., Reich, P. B., Steege, H. t., Morgan, H. D., Heijden, M. G. A. v. d., Pausas, J. G., & Poorter, H. (2003). A handbook of protocols for standardised and easy measurement of plant functional traits worldwide. Australian Journal of Botany, 51(4), 335–380. https://doi.org/10.1071/BT02124

15. Crutsinger, G. M., Collins, M. D., Fordyce, J. A., Gompert, Z., Nice, C. C., & Sanders, N. J. (2006). Plant genotypic diversity predicts community structure and governs an ecosystem process. Science, 313(5789), 966–968. https://doi.org/10.1126/science.1128326

16. Delgado-Baquerizo, M., Maestre, F. T., Reich, P. B., Jeffries, T. C., Gaitan, J. J., Encinar, D., Berdugo, M., Campbell, C. D., & Singh, B. K. (2016). Microbial diversity drives multifunctionality in terrestrial ecosystems. Nature Communications, 7, 10541, Article 10541. https://doi.org/10.1038/ncomms10541

17. Des Roches, S., Post, D. M., Turley, N. E., Bailey, J. K., Hendry, A. P., Kinnison, M. T., Schweitzer, J. A., & Palkovacs, E. P. (2018). The ecological importance of intraspecific variation. Nature Ecology & Evolution, 2(1), 57–64. https://doi.org/10.1038/s41559-017-0402-5

18. Díaz, S., & Cabido, M. (2001). Vive la différence: plant functional diversity matters to ecosystem processes. Trends in Ecology & Evolution, 16(11), 646–655. https://doi.org/10.1016/S0169-5347(01)02283-2

19. Díaz, S., Lavorel, S., de Bello, F., Quetier, F., Grigulis, K., & Robson, M. (2007). Incorporating plant functional diversity effects in ecosystem service assessments. Proceedings of the National Academy of Sciences of the United States of America, 104(52), 20684–20689. https://doi.org/10.1073/pnas.0704716104

20. Diaz, S., Settele, J., Brondizio, E. S., Ngo, H. T., Agard, J., Arneth, A., Balvanera, P., Brauman, K. A., Butchart, S. H. M., Chan, K. M. A., Garibaldi, L. A., Ichii, K., Liu, J., Subramanian, S. M., Midgley, G. F., Miloslavich, P., Molnar, Z., Obura, D., Pfaff, A., Polasky, S., Purvis, A., Razzaque, J., Reyers, B., Chowdhury, R. R., Shin, Y.-J., Visseren-Hamakers, I., Willis, K. J., & Zayas, C. N. (2019). Pervasive human-driven decline of life on Earth points to the need for transformative change. Science, 366(6471), 1327, Article eaax3100. https://doi.org/10.1126/science.aax3100

21. Duffy, J. E. (2003). Biodiversity loss, trophic skew and ecosystem functioning. Ecology Letters, 6(8), 680–687. https://doi.org/10.1046/j.1461-0248.2003.00494.x

22. Eisenhauer, N. (2012). Aboveground-belowground interactions as a source of complementarity effects in biodiversity experiments. Plant and Soil, 351(1-2), 1–22. https://doi.org/10.1007/s11104-011-1027-0

23. Fischer, D. G., Wimp, G. M., Hersch-Green, E., Bangert, R. K., Leroy, C. J., Bailey, J. K., Schweitzer, J. A., Dirks, C., Hart, S. C., Allan, G. J., & Whitham, T. G. (2017). Tree genetics strongly affect forest productivity, but intraspecific diversity-productivity relationships do not. Functional Ecology, 31(2), 520–529. https://doi.org/10.1111/1365-2435.12733

24. George, W., Snecdecor, W. G. C. (1989). Statistical Methods (8th ed.). Iowa State University Press.

25. Grace, J. B. (2006). Structural Equation Modeling and Natural Systems. Cambridge University Press. https://doi.org/DOI: 10.1017/CBO9780511617799

26. Hahn, C. Z., Niklaus, P. A., Bruelheide, H., Michalski, S. G., Shi, M., Yang, X., Zeng, X., Fischer, M., & Durka, W. (2017). Opposing intraspecific vs. interspecific diversity effects on herbivory and growth in subtropical experimental tree assemblages. Journal of Plant Ecology, 10(1), 242–251. https://doi.org/10.1093/jpe/rtw098

27. Hector, A., Schmid, B., Beierkuhnlein, C., Caldeira, M. C., Diemer, M., Dimitrakopoulos, P. G., Finn, J. A., Freitas, H., Giller, P. S., Good, J., Harris, R., Hogberg, P., Huss-Danell, K., Joshi, J., Jumpponen, A., Korner, C., Leadley, P. W., Loreau, M., Minns, A., Mulder, C. P. H., O’Donovan, G., Otway, S. J., Pereira, J. S., Prinz, A., Read, D. J., Scherer-Lorenzen, M., Schulze, E. D., Siamantziouras, A. S. D., Spehn, E. M., Terry, A. C., Troumbis, A. Y., Woodward, F. I., Yachi, S., & Lawton, J. H. (1999). Plant diversity and productivity experiments in European grasslands. Science, 286(5442), 1123–1127. https://doi.org/10.1126/science.286.5442.1123

28. Hillebrand, H., & Matthiessen, B. (2009). Biodiversity in a complex world: consolidation and progress in functional biodiversity research. Ecology Letters, 12(12), 1405–1419. https://doi.org/10.1111/j.1461-0248.2009.01388.x

29. Huang, Y., Chen, Y., Castro-Izaguirre, N., Baruffol, M., Brezzi, M., Lang, A. N., Li, Y., Hardtle, W., Oheimb, G., Yang, X., Liu, X., Pei, K., Both, S., Yang, B., Eichenberg, D., Assmann, T., Bauhus, J., Behrens, T., Buscot, F., Chen, X. Y., Chesters, D., Ding, B. Y., Durka, W., Erfmeier, A., Fang, J., Fischer, M., Guo, L. D., Guo, D., Gutknecht, J. L. M., He, J. S., He, C. L., Hector, A., Hoenig, L., Hu, R. Y., Klein, A. M., Kuehn, P., Liang, Y., Li, S., Michalski, S., Scherer- Lorenzen, M., Schmidt, K., Scholten, T., Schuldt, A., Shi, X., Tan, M. Z., Tang, Z., Trogisch, S., Wang, Z., Welk, E., Wirth, C., Wubet, T., Xiang, W., Yu, M., Yu, X. D., Zhang, J., Zhang, S., Zhang, N., Zhou, H. Z., Zhu, C. D., Zhu, L., Bruelheide, H., Ma, K., Niklaus, P. A., & Schmid, B. (2018). Impacts of species richness on productivity in a large-scale subtropical forest experiment. Science, 362(6410), 80–83. https://doi.org/10.1126/science.aat6405

30. Jactel, H., & Brockerhoff, E. G. (2007). Tree diversity reduces herbivory by forest insects. Ecology Letters, 10(9), 835–848. https://doi.org/10.1111/j.1461-0248.2007.01073.x

31. Johnson, M. T. J., Bertrand, J. A., & Turcotte, M. M. (2016). Precision and accuracy in quantifying herbivory. Ecological Entomology, 41(1), 112–121. https://doi.org/10.1111/een.12280

32. Kahmen, A., Renker, C., Unsicker, S. B., & Buchmann, N. (2006). Niche complementarity for nitrogen: An explanation for the biodiversity and ecosystem functioning relationship? Ecology, 87(5), 1244–1255. https://doi.org/10.1890/0012-9658(2006)87[1244:Ncfnae]2.0.Co;2

33. Kaye, J. P., & Hart, S. C. (1997). Competition for nitrogen between plants and soil microorganisms. Trends in Ecology & Evolution, 12(4), 139–143. https://doi.org/10.1016/s0169-5347(97)01001-x

34. Koricheva, J., & Hayes, D. (2018). The relative importance of plant intraspecific diversity in structuring arthropod communities: A meta-analysis. Functional Ecology, 32(7), 1704–1717. https://doi.org/10.1111/1365-2435.13062

35. Kotowska, A. M., Cahill, J. F., & Keddie, B. A. (2010). Plant genetic diversity yields increased plant productivity and herbivore performance. Journal of Ecology, 98(1), 237–245. https://doi.org/10.1111/j.1365-2745.2009.01606.x

36. Kuznetsova, A., Brockhoff, P. B., & Christensen, R. H. B. (2017). lmerTest Package: Tests in Linear Mixed Effects Models. Journal of Statistical Software, 82(13), 1–26. https://doi.org/10.18637/jss.v082.i13

37. Laforest-Lapointe, I., Paquette, A., Messier, C., & Kembel, S. W. (2017). Leaf bacterial diversity mediates plant diversity and ecosystem function relationships. Nature, 546(7656), 145–147. https://doi.org/10.1038/nature22399

38. Laliberté, E., & Legendre, P. (2010). A distance-based framework for measuring functional diversity from multiple traits. Ecology, 91(1), 299–305. https://doi.org/10.1890/08-2244.1

39. Laliberté, E., Legendre, P., & Shipley, B. (2014). FD: measuring functional diversity from multiple traits, and other tools for functional ecology.

40. Lamb, D., Erskine, P. D., & Parrotta, J. A. (2005). Restoration of degraded tropical forest landscapes. Science, 310(5754), 1628–1632. https://doi.org/10.1126/science.111177

41. Lefcheck, J. S. (2016). PIECEWISESEM: Piecewise structural equation modelling in R for ecology, evolution, and systematics. Methods in Ecology and Evolution, 7(5), 573–579. https://doi.org/10.1111/2041-210x.12512

42. Marquard, E., Weigelt, A., Temperton, V. M., Roscher, C., Schumacher, J., Buchmann, N., Fischer, M., Weisser, W. W., & Schmid, B. (2009). Plant species richness and functional composition drive overyielding in a six-year grassland experiment. Ecology, 90(12), 3290–3302. https://doi.org/10.1890/09-0069.1

43. Müller, M., Klein, A.-M., Scherer-Lorenzen, M., Nock, C. A., & Staab, M. (2018). Tree genetic diversity increases arthropod diversity in willow short rotation coppice. Biomass and Bioenergy, 108, 338–344. https://doi.org/10.1016/j.biombioe.2017.12.001

44. Nabity, P. D., Zavala, J. A., & DeLucia, E. H. (2009). Indirect suppression of photosynthesis on individual leaves by arthropod herbivory. Annals of Botany, 103(4), 655–663. https://doi.org/10.1093/aob/mcn127

45. Nilsson, R. H., Larsson, K.-H., Taylor, A. F S., Bengtsson-Palme, J., Jeppesen, T. S., Schigel, D., Kennedy, P., Picard, K., Glöckner, F. O., Tedersoo, L., Saar, I., Kõljalg, U., & Abarenkov, K. (2019). The UNITE database for molecular identification of fungi: handling dark taxa and parallel taxonomic classifications. Nucleic Acids Research, 47(D1), D259–D264. https://doi.org/10.1093/nar/gky1022

46. Oksanen, a., Blanchet, F. G., Friendly, M., Kindt, R., Legendre, P., McGlinn, D., Minchin, P. R., O’Hara, R. B., Simpson, G. L., Solymos, P., Stevens, M. H. H., Szoecs, E., & Wagner, H. (2019). vegan: Community Ecology Package. https://CRAN.R-project.org/package=vegan

47. Rao, C. R. (1982). Diversity and dissimilarity coefficients: A unified approach. Theoretical Population Biology, 21(1), 24–43. https://doi.org/10.1016/0040-5809(82)90004-1

48. Scherber, C., Eisenhauer, N., Weisser, W. W., Schmid, B., Voigt, W., Fischer, M., Schulze, E. D., Roscher, C., Weigelt, A., Allan, E., Bessler, H., Bonkowski, M., Buchmann, N., Buscot, F., Clement, L. W., Ebeling, A., Engels, C., Halle, S., Kertscher, I., Klein, A. M., Koller, R., Konig, S., Kowalski, E., Kummer, V., Kuu, A., Lange, M., Lauterbach, D., Middelhoff, C., Migunova, V. D., Milcu, A., Muller, R., Partsch, S., Petermann, J. S., Renker, C., Rottstock, T., Sabais, A., Scheu, S., Schumacher, J., Temperton, V. M., & Tscharntke, T. (2010). Bottom- up effects of plant diversity on multitrophic interactions in a biodiversity experiment. Nature, 468(7323), 553–556. https://doi.org/10.1038/nature09492

49. Schmid, B. (1994). Effects of genetic diversity in experimental stands of Solidago Altissima -- evidence for the potential role of pathogens as selective agents in plant populations. Journal of Ecology, 82(1), 165–175. https://doi.org/10.2307/2261395

50. Schöb, C., Kerle, S., Karley, A. J., Morcillo, L., Pakeman, R. J., Newton, A. C., & Brooker, R. W. (2015). Intraspecific genetic diversity and composition modify species-level diversity- productivity relationships. New Phytologist, 205(2), 720–730. https://doi.org/10.1111/nph.13043

51. Schuldt, A., Assmann, T., Brezzi, M., Buscot, F., Eichenberg, D., Gutknecht, J., Haerdtle, W., He, J.-S., Klein, A.-M., Kuehn, P., Liu, X., Ma, K., Niklaus, P. A., Pietsch, K. A., Purahong, W., Scherer-Lorenzen, M., Schmid, B., Scholten, T., Staab, M., Tang, Z., Trogisch, S., von Oheimb, G., Wirth, C., Wubet, T., Zhu, C.-D., & Bruelheide, H. (2018). Biodiversity across trophic levels drives multifunctionality in highly diverse forests. Nature Communications, 9, 2989. https://doi.org/10.1038/s41467-018-05421-z

52. Schuldt, A., Bruelheide, H., Härdtle, W., Assmann, T., Li, Y., Ma, K., von Oheimb, G., & Zhang, J. (2015). Early positive effects of tree species richness on herbivory in a large-scale forest biodiversity experiment influence tree growth. Journal of Ecology, 103(3), 563–571. https://doi.org/10.1111/1365-2745.12396

53. Schuldt, A., Ebeling, A., Kunz, M., Staab, M., Guimarães-Steinicke, C., Bachmann, D., Buchmann, N., Durka, W., Fichtner, A., Fornoff, F., Härdtle, W., Hertzog, L. R., Klein, A.-M., Roscher, C., Schaller, J., von Oheimb, G., Weigelt, A., Weisser, W., Wirth, C., Zhang, J., Bruelheide, H., & Eisenhauer, N. (2019). Multiple plant diversity components drive consumer communities across ecosystems. Nature Communications, 10(1), 1460. https://doi.org/10.1038/s41467-019-09448-8

54. Semchenko, M., Leff, J. W., Lozano, Y. M., Saar, S., Davison, J., Wilkinson, A., Jackson, B. G., Pritchard, W. J., De Long, J. R., Oakley, S., Mason, K. E., Ostle, N. J., Baggs, E. M., Johnson, D., Fierer, N., & Bardgett, R. D. (2018). Fungal diversity regulates plant-soil feedbacks in temperate grassland. Science Advances, 4(11), eaau4578. https://doi.org/10.1126/sciadv.aau4578

55. Siefert, A., Violle, C., Chalmandrier, L., Albert, C. H., Taudiere, A., Fajardo, A., Aarssen, L. W., Baraloto, C., Carlucci, M. B., Cianciaruso, M. V., de L. Dantas, V., de Bello, F., Duarte, L. D. S., Fonseca, C. R., Freschet, G. T., Gaucherand, S., Gross, N., Hikosaka, K., Jackson, B., Jung, V., Kamiyama, C., Katabuchi, M., Kembel, S. W., Kichenin, E., Kraft, N. J. B., Lagerström, A., Bagousse-Pinguet, Y. L., Li, Y., Mason, N., Messier, J., Nakashizuka, T., Overton, J. M., Peltzer, D. A., Pérez-Ramos, I. M., Pillar, V. D., Prentice, H. C., Richardson, S., Sasaki, T., Schamp, B. S., Schöb, C., Shipley, B., Sundqvist, M., Sykes, M. T., Vandewalle, M., & Wardle, D. A. (2015). A global meta-analysis of the relative extent of intraspecific trait variation in plant communities. Ecology Letters, 18(12), 1406–1419. https://doi.org/10.1111/ele.12508

56. Tilman, D., Reich, P. B., Knops, J., Wedin, D., Mielke, T., & Lehman, C. (2001). Diversity and productivity in a long-term grassland experiment. Science, 294(5543), 843–845. https://doi.org/10.1126/science.1060391

57. Turnbull, L. A., Isbell, F., Purves, D. W., Loreau, M., & Hector, A. (2016). Understanding the value of plant diversity for ecosystem functioning through niche theory. Proceedings of the Royal Society B: Biological Sciences, 283(1844), 20160536. https://doi.org/10.1098/rspb.2016.0536

58. van der Heijden, M. G. A., Bardgett, R. D., & van Straalen, N. M. (2008). The unseen majority: soil microbes as drivers of plant diversity and productivity in terrestrial ecosystems. Ecology Letters, 11(3), 296–310. https://doi.org/10.1111/j.1461-0248.2007.01139.x

59. van Moorsel, S. J., Hahl, T., Wagg, C., De Deyn, G. B., Flynn, D. F. B., Zuppinger-Dingley, D., & Schmid, B. (2018). Community evolution increases plant productivity at low diversity. Ecology Letters, 21(1), 128–137. https://doi.org/10.1111/ele.12879

60. van Moorsel, S. J., Schmid, M. W., Wagemaker, N. C. A. M., van Gurp, T., Schmid, B., & Vergeer, P. (2019). Evidence for rapid evolution in a grassland biodiversity experiment. Molecular Ecology, 28(17), 4097–4117. https://doi.org/10.1111/mec.15191

61. Vellend, M., & Geber, M. A. (2005). Connections between species diversity and genetic diversity. Ecology Letters, 8(7), 767–781. https://doi.org/10.1111/j.1461-0248.2005.00775.x

62. Williams, L. J., Paquette, A., Cavender-Bares, J., Messier, C., & Reich, P. B. (2017). Spatial complementarity in tree crowns explains overyielding in species mixtures. Nature Ecology & Evolution, 1(4), 0063. https://doi.org/10.1038/s41559-016-0063

63. Zuppinger-Dingley, D., Schmid, B., Petermann, J. S., Yadav, V., De Deyn, G. B., & Flynn, D. F. B. (2014). Selection for niche differentiation in plant communities increases biodiversity effects. Nature, 515(7525), 108–111. https://doi.org/10.1038/nature13869

64. Zvereva, E. L., Zverev, V., & Kozlov, M. V. (2012). Little strokes fell great oaks: minor but chronic herbivory substantially reduces birch growth. Oikos, 121(12), 2036–2043. https://doi.org/10.1111/j.1600-0706.2012.20688.x

